# Genetic buffering and potentiation in metabolism

**DOI:** 10.1101/845792

**Authors:** Juan F. Poyatos

## Abstract

Cells adjust their metabolism in response to mutations, but how this reprogramming depends on the genetic context is not well known. Specifically, the absence of individual enzymes can affect reprogramming and thus the impact of mutations in cell growth. Here, we examine this issue with an *in silico* model of *Saccharomyces cerevisiae’s* metabolism. By quantifying the variability in the growth rate of 10000 different mutant metabolisms that accumulated changes in their reaction fluxes, in the presence, or absence, of a specific enzyme, we distinguish a subset of modifier genes serving as buffers or potentiators of variability. We notice that the most potent modifiers refer to the glycolysis pathway and that, more broadly, they show strong pleiotropy and epistasis. Moreover, the evidence that this subset depends on the specific growing condition strengthens its systemic underpinning, a feature only observed before in a simple model of a gene-regulatory network. Some of these enzymes also modulate the effect that biochemical noise and environmental fluctuations produce in growth. Thus, the reorganization of metabolism triggered by mutations has not only direct physiological implications but also changes the influence that other mutations have on growth. This is a general result with implications in the development of cancer therapies based on metabolic inhibitors.

## Introduction

Cells experience mutations in different ways, and the direct importance of these on the phenotype has been the focus of substantial basic and applied research (Nagy et al., 2003; Stratton et al., 2009). It is much less known, however, how specific genetic contexts modify the phenotypic impact of mutations (Chow, 2016; Dowell et al., 2010) and the many consequences that the alterations could have in disease progression (Ashworth et al., 2011).

One can expect two broad situations. In the first one, the presence of particular genetic variants buffers the effect of mutations. This result helps explain the robustness observed in biological phenotypes and was already discussed –under the notion of canalization– in early studies of development (Rendel, 1967; Schmalhausen, 1949; Waddington, 2014). Canalization, or robustness, also leads to the accumulation of cryptic genetic variation (Gibson and Dworkin, 2004; Paaby and Rockman, 2014) that is normally hidden under typical conditions. Thus, the unveiling of this hidden variation after perturbation was considered to be a measure of a decline of robustness. However, this is not necessarily so (Hermisson and Wagner, 2004; Richardson et al., 2013): two systems presenting the same robustness can nevertheless expose cryptic variation linked to mutations which are neutral depending on the system they emerge (conditional neutrality)(Hermisson and Wagner, 2004; Paaby and Rockman, 2014; Richardson et al., 2013). Moreover, a second general scenario corresponds to the case in which some genetic variants potentiate the functional consequences of mutations what can eventually promote the rapid evolution of new traits (Cowen and Lindquist, 2005; Whitesell et al., 2014).

This wide range of implications encouraged the search for the genetic underpinning of buffering or potentiation. And thus, the chaperone Hsp90 was the first described protein deemed to be a canalization agent, initially demonstrated in *Drosophila* (Rutherford and Lindquist, 1998) and later generalized across species (Jarosz and Lindquist, 2010; Queitsch et al., 2002; Rohner et al., 2013). Hsp90 represents a buffer, or capacitor, whose effects in the folding and stability of other proteins fits well with the notion of a global element contributing to the canalized phenotype, a role that has also been attributed to a few additional molecular agents, like the prion [PSI^+^](Tyedmers et al., 2008).

But later studies raised some doubts on the action, definition, and uniqueness of certain proteins as capacitors. For instance, in the precise case of heat shock proteins, part of the associated variation is linked to their control of mutagenic activity by transposons (Specchia et al., 2010). Besides, these proteins can not only reduce but also amplify the impact of mutations by letting them have immediate phenotypic consequences. The same molecular element is therefore modifying the impact of mutations in two contrasting ways (Cowen and Lindquist, 2005; Whitesell et al., 2014). Other uncertainties indicate constraints on the conventional experimental approach to examine these issues, in which selection sieves the mutations commonly assayed. Mutation accumulation experiments (Halligan and Keightley, 2009) reduce the strength of selection and thus provide a more accurate sample instead (Geiler-Samerotte et al., 2018).

Beyond these objections, a more important criticism is the evidence that buffers, or potentiators, are not fundamentally connected to special molecular agents with distinct biochemical properties but that they emerge as an intrinsic feature of complex biological networks. Many genes could then modify the effect of mutations (Bergman and Siegal, 2003); a prominent conclusion if one were to bring in the earlier results as part of the representative methodology of genetics (Nagy et al., 2003) but maybe less unexpected in the broader framework of the architecture of complexity (Simon, 1962).

In this manuscript, we consider metabolism as a representative model system to examine whether buffering and potentiation is indeed a common phenomenon in biological networks. While this result has been shown with the use of simple gene-regulatory networks (Bergman and Siegal, 2003), and the finding of new genes acting as capacitors further confirms this conclusion (Richardson et al., 2013), its validation in more realistic networks is still lacking. Moreover, and given that the experimental manipulations accompanying this question are challenging, we contemplate instead an *in silico* representation of metabolism, with the advantage of enabling a full mechanistic account of the phenomena considered. Earlier work on robustness and evolution of metabolic networks confirms the soundness of this approach (Barve et al., 2012; Ho and Zhang, 2016; Pál et al., 2005).

We thus consider a genome-scale reconstruction of *Saccharomyces cerevisiae*(Duarte, 2004) to ask if the presence of a particular enzyme could change the influence on growth rate of a compendium of mutations altering the metabolic fluxes. To this aim, we generate a collection of mutant metabolisms (mutation accumulation lines) derived from the wild-type, which displays a distinct growth rate distribution. We then quantify if these *very same* lines change the growth rate in a different manner depending on the absence of a single enzyme. This led us to identify a set of genes acting as buffers and potentiators whose influence depends on the particular working conditions (i.e., type of available nutrients) of the metabolism, and the sources of variability considered. We finish analyzing how this fundamental phenomenon could have practical implications in the development of metabolic-based cancer therapies.

## Results

### Buffers and potentiators in metabolism

We examined the significance of each metabolic enzyme on how mutations impact growth rate, which is regarded here as a case study of a complex phenotype. To this aim, we generated a collection of mutant metabolisms that simulates the production of spontaneous mutations in independent cell lines, like those obtained in mutation-accumulation (MA) experiments (Halligan and Keightley, 2009). This kind of collections helps characterize the response of biological systems to new mutations which did not experience any purge by selection (Hermisson and Wagner, 2004).

In this metabolic setting, we first derived the mutant compendium by limiting the flux of 5% of the total reactions chosen randomly in the wild-type metabolism (**Figure 1A**). We obtained in this way 10000 different mutant lines, a feasible number to generate *in silico*, but a challenging one to reproduce experimentally (a typical MA collection contains about 100 lines (Halligan and Keightley, 2009)). For each member of the compendium, we compute its growth rate (“fitness”) by minimizing the metabolic adjustment caused by the mutations on the fluxes of the wild-type metabolism, an approach that is known to successfully predict growth rates and fluxes upon mutation (Segre et al., 2002). Each line included in the collection presents nonzero fitness (see Methods for details).

**Figure 1.**
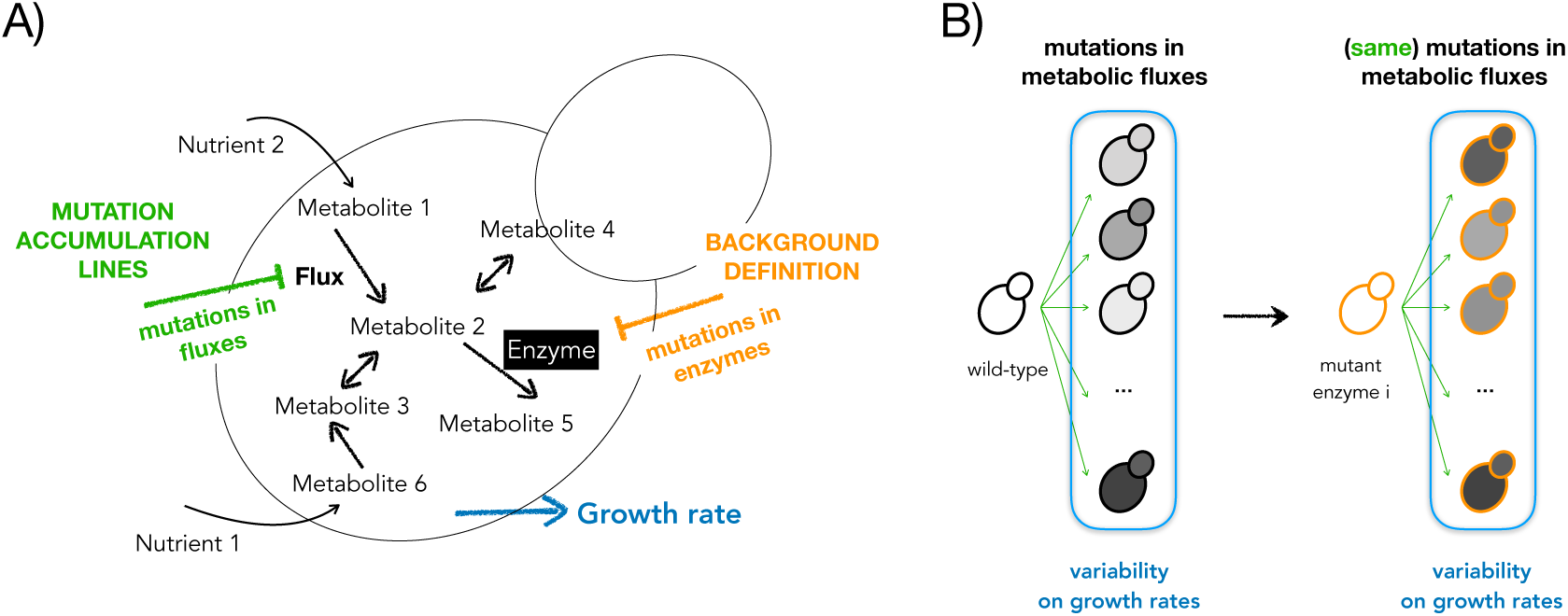
Influence of global modifiers in metabolism. A) An *in silico* representation of yeast metabolism can experience two types of mutations: 1)mutations in metabolic fluxes, which define the mutation accumulation lines, and 2)mutations in the enzymes, which define the particular backgrounds. The complex phenotype considered is growth rate (relative to the corresponding growth rate of each reference metabolism). B) We score the variability of the growth rates in a group of different lines (arrows) in which mutations in the metabolic fluxes are accumulated. We compute this variability in the presence (wild-type) and the absence (mutant metabolism) of a particular enzyme i. Here the difference between growth rates and metabolic backgrounds are represented by the colors of the fill and the border of the yeast cartoons, respectively.

We then computed the relative effect of the preceding MA lines in any metabolism in which an individual enzyme has been deleted, i.e., the mutations constituting the MA lines are fixed (**Figure 1B**). The difference in phenotypic (growth-rate) variation in the presence or absence of an enzyme reveals how it modifies the consequences of flux mutations on fitness. We quantified this difference with a score defined by the change between standard deviations θ = (std_mutant_ – std_wild-type_)/std_wild-type_, with θ < 0 indicating that the enzyme works as a potentiator (presence of the enzyme increases variability) and θ > 0 indicating that it acts as a buffer [presence of the enzyme decreases variability, Methods (Geiler-Samerotte et al., 2018; Hermisson and Wagner, 2004)].

Under a nutrient-rich condition (YPD), and after filtering out enzymes with no effect and isoenzymes, we identified 14 enzymes that significantly modify the response to mutations (**Figure 2, Table S1**, Methods). Within this set, we also recognized five cases of particularly strong effects, which are all related to the glycolysis/gluconeogenesis pathway: PGK1 (potentiator) and TPI1, PGI1, FBA1 and PFK1 (buffers). Deletion of these enzymes lead to particularly strong flux rewiring (observed rewiring = 76% of the total flux in the wild-type, mean rewiring expected randomly = 2%, random permutation test, p < 1e-4, with 10000 permutations) low fitness of the associated mutated metabolism (observed relative fitness by FBA = 0.2, mean relative fitness expected randomly = 0.98, random permutation test as before, p < 1e-4) and increase number of MA lines with no growth (observed number of lethal MA lines = 1016, mean number of lethal lines expected randomly = 36, random permutation test as before, p < 1e-4; total number of lines = 10000); all features denoting the occurrence of very strong metabolic readjustments due to the enzyme deletion (Methods).

**Figure 2.**
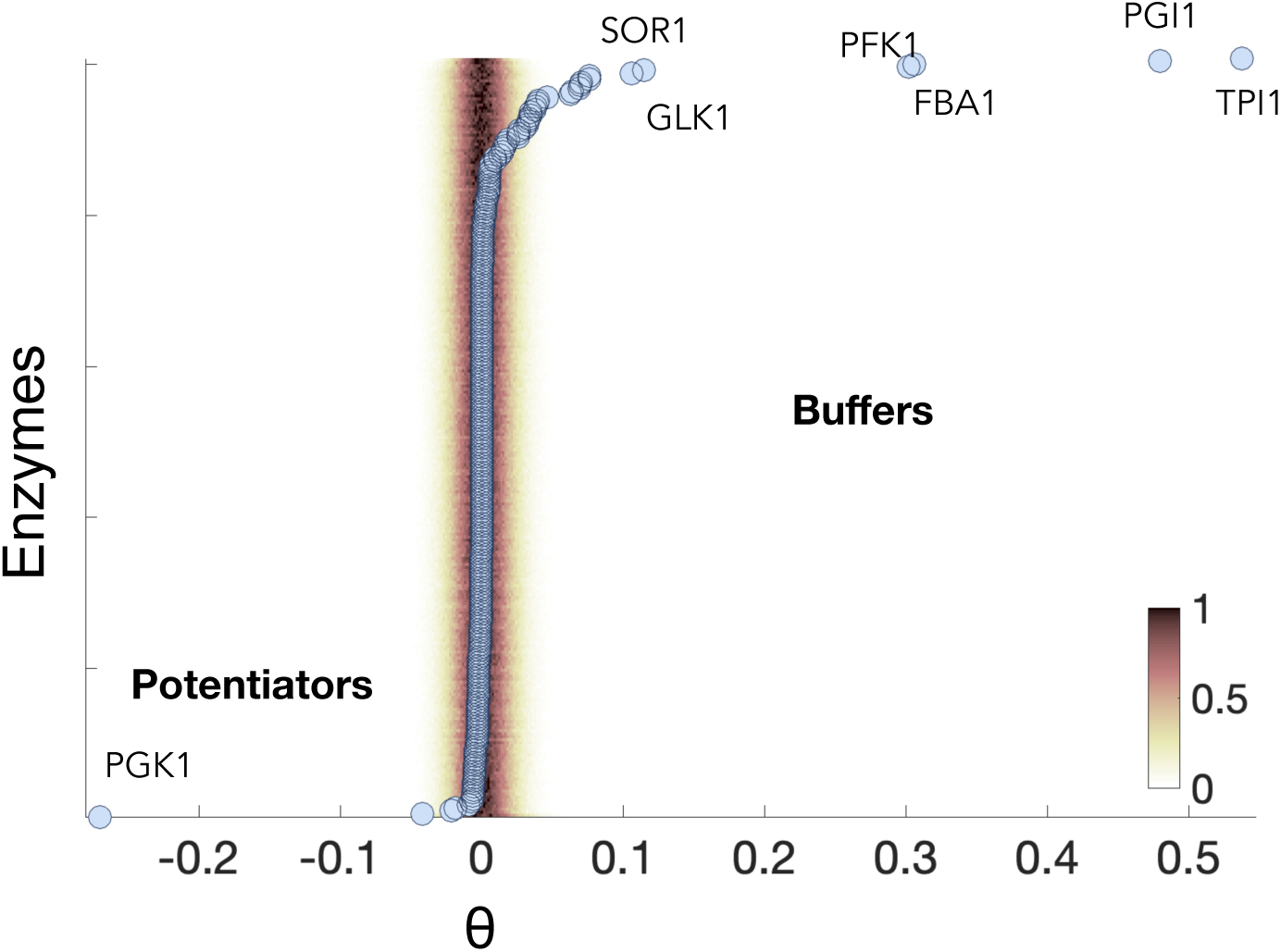
Buffers and Potentiators in yeast metabolism. For each enzyme, a θ score is computed, which is proportional to the difference in variability between the mutant and wild-type backgrounds. The shadow denotes the normalized null probability distribution of getting a particular score for each metabolic background. We added the names of the most significant modifiers (buffers with θ>0 and potentiators with θ<0).

### Metabolic rationale underlying buffers and potentiators

The advantage of *in silico* models is that these readjustments can be uncovered. Thus, an enzyme works as a potentiator when its absence frequently disables the costs of mutating a significant number of reactions, included in the MA lines, which decreases variability in growth rate (std_mutant_ < std_wild-type_). To evaluate this, we identified those reactions enriched in MA lines whose impact on growth rate decreased in the ΔPGK1 background. The top five belong to the glycolysis-gluconeogenesis system and pyruvate metabolism. This is reasonable considering that PGK1 (3-PhosphoGlycerate Kinase) is a key enzyme whose mutation inactivates the fluxes on these pathways. The cost in growth of a mutation on these reactions is, therefore, smaller than in the wild-type background.

Enzymes working as buffers have the opposing effect. In this case, the absence of a buffer implies that the weight of a substantial number of mutations within the MA lines is amplified, so that the overall variability in growth increases (std_mutant_ > std_wild-type_). Which type of mutations show this amplification depends again on the effect of the specific background. If we first consider the top four buffers with strong effect, we identify several reactions that considerably increased the flux in the corresponding metabolic background, like those related to alternative carbon metabolisms, e.g., glycerol, sorbitol, etc.

Beyond the specifics of the metabolic readjustments, both pleiotropy and epistasis have been argued to be relevant features to interpret buffers and potentiators. They quantify the functional role and number of interactions with other mutations of these elements, respectively (Richardson et al., 2013). We consequently examined both features by computing the epistatic network between every pair of enzymes (Segrè et al., 2005) [but note that higher order interactions are also important (Kuzmin et al., 2018; Taylor and Ehrenreich, 2014)] and a metabolic pleiotropic score recently introduced that quantifies the contribution to each enzyme to every biomass precursor (Shlomi et al., 2007; Szappanos et al., 2011). Global modifiers show strong pleiotropy and epistasis (**Table S1**). This indicates overall their multifunctionality character as illustrated in **Figure 3** where pleiotropy and the number of weak negative genetic interactions are explicitly shown. Note that this specific class of genetic interactions reflects a multiplicity of alternatives to perform a specific function (Bajić et al., 2014) (Methods).

**Figure 3.**
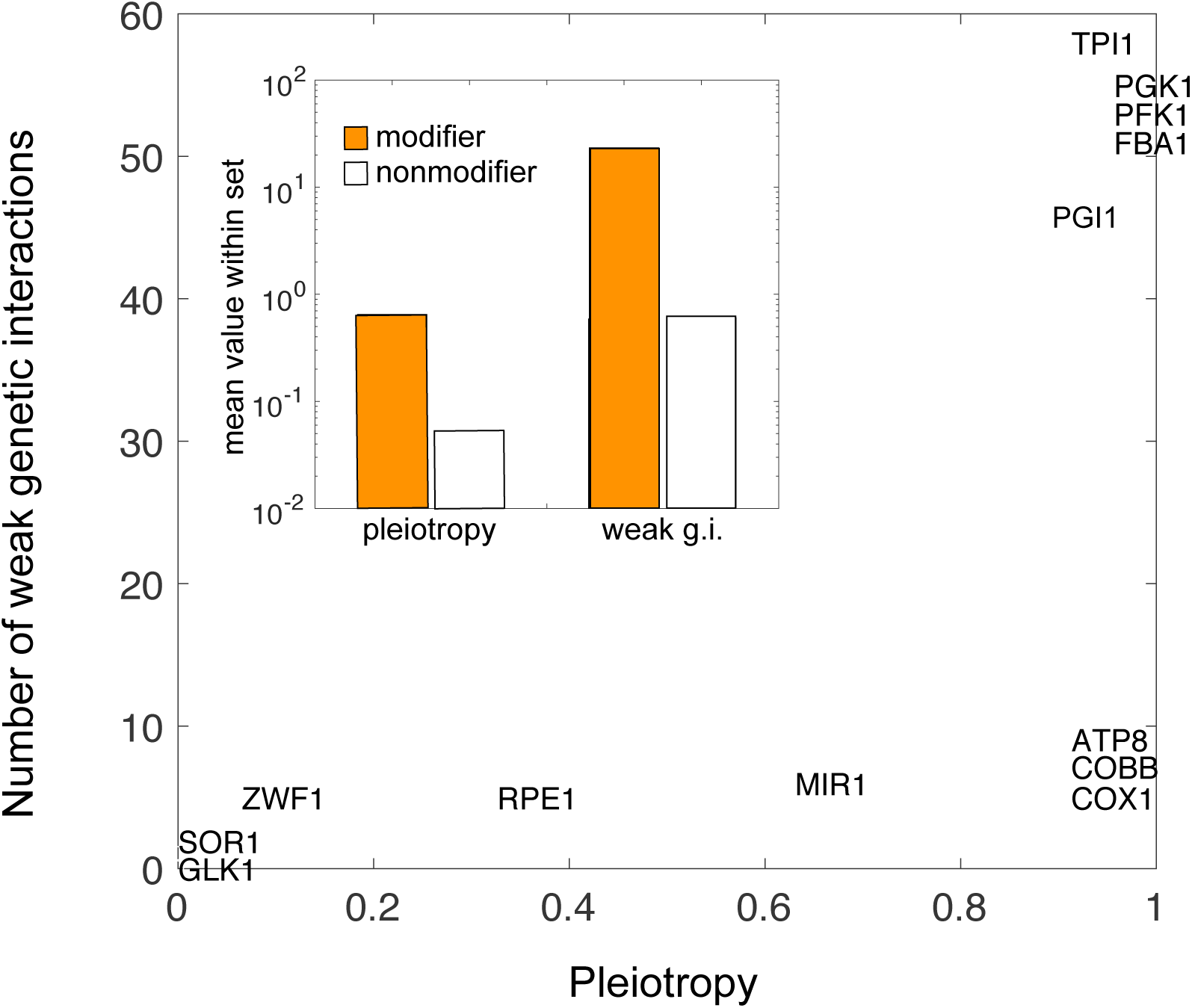
Buffers and potentiators correspond to multifunctional enzymes. We computed the pleiotropy and number of weak negative genetic interactions as proxies of enzyme multifunctionality (see main text and Methods). Modifiers (both buffers and potentiator) show stronger pleiotropy (mean pleiotropy modifiers = 0.65, mean pleiotropy nonmodifiers = 0.05, two-sample Kolmogorov-Smirnov test p = 1.6e-6) and number of weak negative genetic interactions (g.i.) (mean number of weak g.i. modifiers = 23.38, weak g.i. nonmodifiers = 0.6, KS p = 6.9e-8).

### Buffers and capacitors are condition dependent

These results confirm the intrinsic presence of buffering and potentiation elements modulating the response to mutations in biological networks, a result only discussed before with the use of simple gene-regulatory network models and that we extend here to a representative metabolic setting. Moreover, and given that the function of metabolic networks strongly depends on the precise growing conditions (Papp et al., 2004), we could expect that most enzymes modify the response to mutations in a condition-dependent manner. To examine this, we studied two complementary situations (Methods), one in which we modify the carbon source complementing YPD conditions (that includes glucose by default), and a second one in which we studied a range of random nutrient conditions (from poor to rich media), and also minimal medium.

As projected, the list of enzymes acting as buffers or potentiators generally changes, with some enzymes precisely related to the specific growing conditions **(Table S2)**. For instance, GAL1, GAL7, and GAL10 (related to galactose metabolism) act as potentiators in YPG medium (galactose as carbon source), while glycerol utilization enzymes (GUT1 and GUT2) are potentiators in YPGly (glycerol as carbon source). Other enzymes change their role, e.g., TPI1 (Triose-Phosphate Isomerase) functions as a buffer when growing in minimal medium or a potentiator in YPGly. In contrast, COX1, COBB, and ATP8 consistently buffer variation. Besides, and while there is a general tendency to exhibit more buffering that potentiation, there exist situations in which potentiation is dominant and others in which the number of enzymes acting as buffers is severely reduced. This emphasizes that the role of a particular enzyme in modifying the impact of mutations is a systemic feature of metabolism that depends on its regime of activity, i.e., a change of environmental condition matters.

### Are there enzymes acting as universal modifiers?

All the previous analyses distinguish a set of genes that can modify the amount of growth rate variability caused by the accumulation of mutations. We were also interested in studying to what extent these enzymes represent “universal” modifiers, i.e., their absence also changes the response to other sources of phenotypic variation. If this were the case, it would suggest a single mode of canalization, i.e., the presence of broad mechanisms to alter the effect of perturbations (Meiklejohn and Hartl, 2002). Note, however, that more recent results argued against this congruence (Dworkin, 2005; Richardson et al., 2013). The debate is still open and surely depends on the level of the biological organization considered. We tried to examine this issue here with regards to two additional sources of variability; biochemical noise (related to the low copy number of molecules) and environmental fluctuations.

To generate the variability coupled to biochemical noise, we need first to compute the noise corresponding to the flux of each metabolic reaction in the network. We followed a previously established approach (Wang and Zhang, 2011) (Methods), which uses data on expression noise of the enzymes (obtained in YPD medium) and explicit knowledge about the metabolic logic to subsequently estimate the reaction noise (Wang and Zhang, 2011). We can then consider 10000 independent realizations in which the flux of metabolic reactions is randomized depending on its noise (Methods). We thus obtain a distribution of growth rates for the wild-type metabolism and for those genetic backgrounds in which each of the enzymes is deleted. This permits us to compute a θ score as previously, but concerning the variability in growth rate due to biochemical noise: θ_noise_.

We noticed that the five strongest modifiers with respect to mutations also appear as modifiers regarding noise; PGK1 as potentiator and PFK1, FBA1, TPI1 and PGI1 as buffers (although PGI1 emerges as a very strong buffer in this case instead of TPI1, the strongest buffer to variability caused by mutations). Three other “mutational” buffers remain as such: SOR1, RPE1, and GLT1, while new enzymes merely buffering variability due to noise also appear: GRE3, MAE1, CTT1, etc. (**Table S1**).

We next examined the response to fluctuations in the environmental conditions (Dworkin, 2005). By this we mean deviations on the import fluxes that characterize YPD. To generate a fitness distribution, we computed growth rate in 10000 different environments in which the import of the corresponding nutrients fluctuates 10% of its fixed YPD value (Methods). Fitness distributions were again computed for the wild-type and for all metabolisms in which one enzyme has been deleted to compute θ_environment_. This θ score is proportional to std_mutant_ – std_wild-type_ as before.

In this setting, we find again that PGK1 acts as a potentiator and that PGI1, COBB, COX1, and ATP8 remain as buffers (**Table S1**). Thus, there are two central enzymes that act as a potentiator (PGK1) or buffer (PGI1) to all three sources of growth rate variability. Moreover, we computed the correlation of all three scores obtained (for every enzyme) as a measure of the similarity in the mode of canalization. We detect the strongest correlation between the mutational and the noise-induced variability (R = 0.77, p = 1.74e-101; mutational and environmental R = 0.45, p = 6.58e-14, noise and environmental, R = 0.35, p = 1.76e-08).

## Discussion

The interconnectedness of biological systems, as revealed by the widespread identification of pleiotropic and epistatic effects (Kuzmin et al., 2018), suggests that the presence of genetic modifiers of phenotypic variability should be a prevalent phenomenon (Hermisson and Wagner, 2004; Waddington, 2014). Gene-regulatory models (Bergman and Siegal, 2003) and morphometrics experiments in *Drosophila* (Takahashi, 2013) appear to confirm such a view. But finding additional contexts to validate this principle is challenging given the insufficiency of large-scale experimental approaches to examine phenotypic variation with high resolution.

Here, we use genome-scale metabolic models to generate large-scale quantitative phenotypic data. We show that many enzymes work as buffers or potentiators of phenotypic variability originated by mutations in the reaction fluxes, with growth rate representing the complex phenotype. This set is dependent on the precise working regime of the metabolism, e.g., the growing medium, emphasizing that this is an intrinsic property of the system generating the phenotype rather than of its constituents. In most of these regimes we detected suppression of variation (buffering) as projected with simpler models (Bergman and Siegal, 2003), but there exist certain conditions in which potentiation predominates.

A particular enzyme might similarly be a modifier for other sources of variability (Meiklejohn and Hartl, 2002). We explicitly studied variability generated by the presence of biochemical noise or fluctuating environmental (nutrient) conditions. We find that congruence is particularly strong between mutational and noise perturbations. However, given that our protocol to generate these two types of variability affects fluxes in a qualitatively similar manner, it is not surprising that we encounter similitude between the corresponding set of modifiers. Metabolism may nevertheless represent a particular biological system in which different perturbations eventually lead to the same response, but some disparities could also be observed.

Moreover, the strongest modifiers we notice, which comprise the main enzymes of the glycolysis and respiratory chain, validate the idea that it is the multi-functionality of these elements within the network that matters. Both sets of enzymes showed strong pleiotropy, which also correlates with the extent of metabolic rewiring and the amount of change of genetic interactions experienced when these enzymes are mutated (Bajić et al., 2014). That we uncover a similar set if we consider a different metabolic model (Methods) validates our exploration. More work on metabolic models would, of course, improve the individual predictions (Heavner and Price, 2015).

Note also that four of the strongest modifiers (PGK1, FBA1, PGI1, and TPF1) are essential genes and that the model also predicts strong fitness costs (**Table S1**). This confirms previous reports presenting essential genes as principal agents in regulating phenotypic variance (Levy and Siegal, 2008). The result was based on morphometric phenotypes, so we were interested to examine if the modifiers we obtained here might also represent modifiers to these additional traits. Variability is summarized in this case by introducing a phenotypic potential’ both in nonessential (Levy and Siegal, 2008) and essential genes (Bauer et al., 2015)]: how much a mutation changes morphological variation. We plot the distribution of these scores in **Figure 4**, and also the precise value corresponding to the (metabolic) modifiers to fitness. Only two of them remain as modifiers.

**Figure 4.**
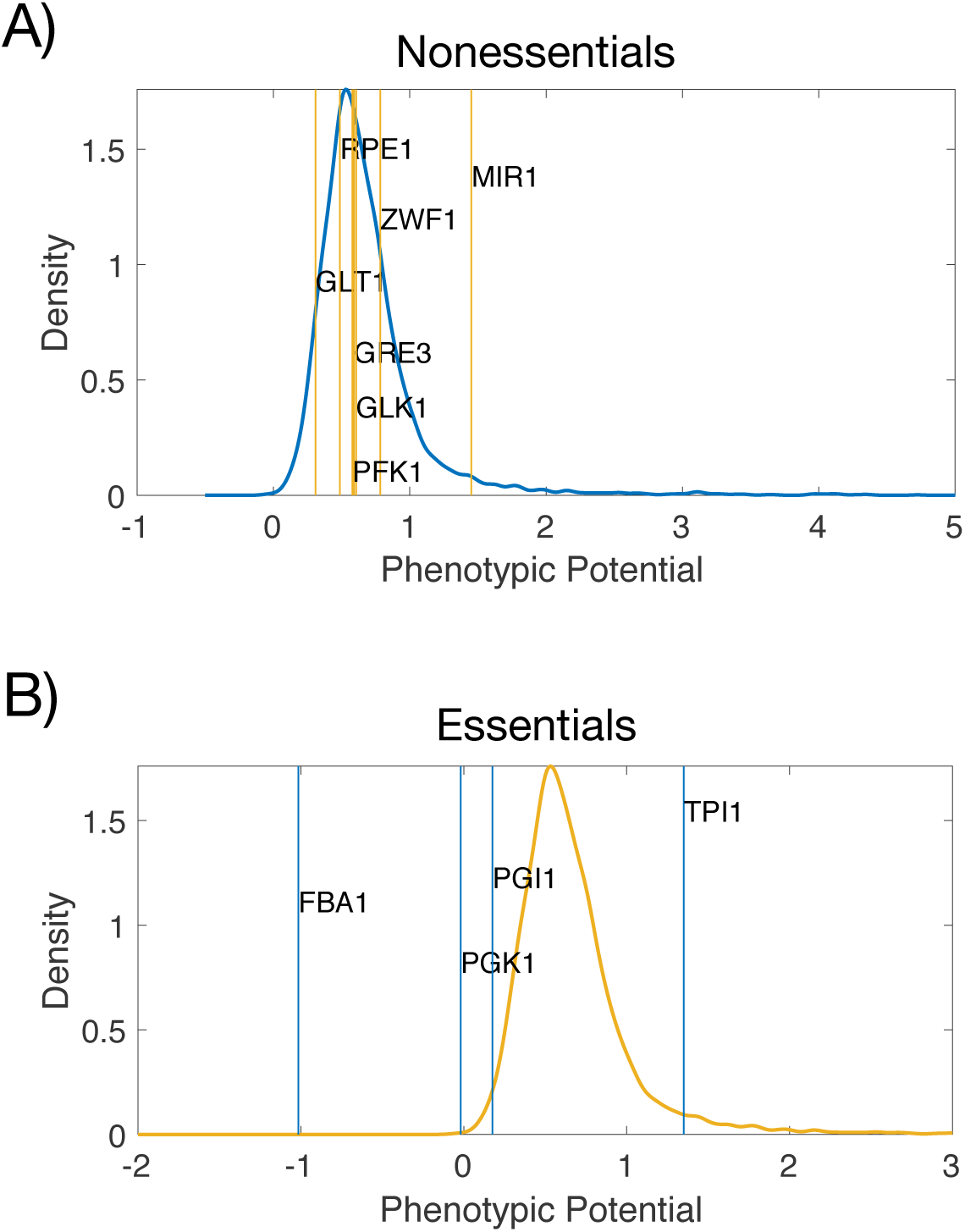
Phenotypic potential to morphological variation. The phenotypic potential scores the amount of morphological variation associated to a specific mutation (Methods). We show a kernel density plot of the distribution of scores for a collection of nonessential (A) and essential (B) genes, and the corresponding values for the set of modifiers to growth rate as phenotype (obtained in YPD; see Figure 2). MIR1 (nonessentials) and FBA1 (essentials) significantly exhibit a larger and smaller phenotypic potential than expected (p < 0.05, randomizing test), respectively.

Finally, our work has connotations for the study of how metabolic heterogeneity occurs in tumors emphasizing its very dynamic nature. Specific acquired mutations cause metabolic reprogramming (e.g., oncogenic drivers leading to characteristic metabolic signatures), but the current knowledge of this rewiring is somehow coarse; one mutation, or combination of mutations, leads to certain rewiring. Other work already hinted, however, to more complicated results associated with the rewiring produced by the same mutation, e.g., influence of tissue of origin, or cell lineage, etc. (DeBerardinis and Chandel, 2016; Hu et al., 2013). We have seen here how mutations influence the fitness effects of added mutations, and how they decisively shape the amount of heterogeneity. Indeed, to partially evaluate this effect, we examined — using data from a collection of tumor samples— how the mutations characterizing particular cancers modify the variation in the expression of metabolic enzymes. We observed that the variation is typically larger within tumor samples that within the corresponding controls, which suggests that the mutated genes contribute to buffering against environmental variability (Methods, **Fig. S1**).

Furthermore, the impact of oncogenic drivers depends on the microenvironment, which could change, again as we have appreciated here, the role of a precise gene mutation as buffer or potentiator of phenotypic variability (Geiler-Samerotte et al., 2016). These feedbacks eventually influence cancer progression and might have consequences in several therapies that are targeting different parts of metabolism, normally the glycolysis pathway. While the consequences of acting on certain targets might depend on the characteristic metabolic reprogramming linked to the genetic lesion and tissue type (Yuneva et al., 2012), this work accentuates that the outcome of metabolic inhibitors goes beyond the alteration of metabolism to the modification of the phenotypic consequences of subsequent mutations. Work is needed to assess the influence of this component in the application of effective interventions (Riordan and Nadeau, 2017) to prevent disease.

## Acknowledgements

I would like to thank Eugene Plavskin and Mark L Siegal for discussions and the Center for Genomics and Systems Biology, New York University, for their hospitality while this research was conducted. I also thank Alvar Alonso-Lavin, Djordje Bajić, Mónica Chagoyen, and Pablo Yubero for comments on an earlier version the manuscript. This work was supported in part through the NYU IT High Performance Computing resources, services, and staff expertise, and grants FIS2016-78781-R and the Salvador de Madariaga program (grant PRX18/00439) from the Spanish Ministerio de Economía y Competitividad.

## Materials and Methods

### Models

We mostly worked with *Saccharomyces cerevisiae* iND750 (Duarte, 2004) with a total number of 1266 reactions that incorporates all necessary complexity from yeast metabolism, while enabling us to moderate the substantial computational load associated with our analysis. Standard conditions correspond to YPD rich medium (with 20 mmol gr^−1^h^−1^ of glucose and 2 mmol gr^−1^h^−1^ of O_2_ import and an assortment of amino acids introduced at a rate of 0.5 mmol gr^−1^h^−1^). Reactions in the model are part of 56 subsystems linked to different metabolisms, e.g., fatty acids, glutamate, etc. Within these subsystems, two corresponds to exchange and biomass reactions (117 reactions) and bicarbonate (HCO_3_) equilibration reactions. To validate the general appearance of buffers and potentiators in metabolism, we examined an additional *Saccharomyces cerevisiae* model [iAZ900 (Zomorrodi and Maranas, 2010), in YPD medium]. Using this model, we identified three potentiators (including PGK1), and sixteen buffers (including, ATP8, RPE1, SOR1, GLK1, COBB, COX1, TPI1, PGI1, FBA1, and PFK1) (**Table S3**).

### FBA and MOMA

FBA is a mathematical tool for metabolic network analysis that allows the prediction of growth rate, i.e., fitness, and fluxes under the assumption of maximization of biomass production given a set of constraints. We use the Gurobi linear programming optimizer (www.gurobi.com) and the Cobra toolbox (Heirendt et al., 2019) in Matlab (www.mathworks.com). We also minimize the absolute value of fluxes to avoid loops in the solutions. We compute all reference metabolisms (wild-type and single-enzyme deletions, see below) with FBA. To obtain the fitness for each of the components of a MA mutation line, after deletions, we used MOMA, a procedure that minimizes the deviation in fluxes from the corresponding metabolism without the mutations. MOMA outperforms the standard FBA approach in the prediction of growth rate and fluxes upon mutation relying on the assumption that after genetic perturbations, the organism’s metabolic and regulatory responses favour a new steady state close to the original operating region, rather than maximizing cellular growth (Segre et al., 2002).

### Generation of mutation accumulation lines

We produced 10000 independent “mutation accumulation lines” by fixing for each line the flux of 5% of the constituent biochemical reactions of the wild-type metabolism chosen at random. For each designated reaction, we assigned a random value obtained from a uniform distribution between 0 and 20 mmol gr^−1^h^−1^ to the corresponding lower (reversible reactions) and upper bounds of the associated flux. Reactions involved in external exchange (116 reactions) are not incorporated in the generation of the MA lines to maintain the nutrient conditions.

### Protocol to identify buffers and potentiators

Our goal is to quantify to what extent the accumulation of a fixed set of mutations (“MA lines”) causes a different response in growth due to the presence or absence of a particular enzyme. We begin with a compilation of “reference” metabolisms that includes the wild-type and all possible variants with a single enzyme removed. The growth rate of these metabolisms is computed with FBA. After this, each reference metabolism experiences the very same set of mutations in the fluxes (the MA lines defined previously). For each line, fitness is calculated with MOMA with regards to deviations to the respective reference metabolism and normalized by the fitness value of the reference (all lines with the wild-type as a reference has nonzero fitness). We calculated the variability on the (relative) fitness observed in the 10000 MA lines. If the variability observed in a specific mutant is bigger than that observed in the wild-type, we say that the corresponding enzyme is a buffer; if smaller, we say that it is a potentiator. We use the relative difference with respect to the wild-type value θ = (STD fit_mutant – STD fit_wild-type)/STD fit_wild-type (Geiler-Samerotte et al., 2018; Hermisson and Wagner, 2004) as score. Different measures that we tested led to comparable results, like the genotype-by-line interaction variance (Richardson et al., 2013) (**Table S1**).

### Flux rewiring

For each reference metabolism, we computed the Euclidean norm of the vector defined by the difference between the optimal FBA fluxes of the mutant background and the wild-type. We divided this value by the Euclidean norm of the optimal wild-type FBA flux. This indicates the degree of metabolic reprogramming experienced by a given mutant.

### Random environments, environments with a carbon source other than glucose and minimal medium

Random environments were aerobic (2 mmol gr^−1^h^−1^ of O_2_ import; ammonia, phosphate, sulphate, sodium, potassium, CO_2_ and H_2_O unbound), with the specific set of nutrients being selected from an exponential distribution probability (Wang and Zhang, 2009) (with mean = 0.1). After defining this set, their dosage was randomly obtained by applying a uniform distribution between 0 and 20 mmol gr^−1^h^−1^ (**Table S2**). We also examined some YPD variants, i.e., YPE, YPGal, YPGly, and YPLac, in which the import of glucose at 20 mmol gr^−1^h^−1^ as a carbon source is substituted by ethanol, galactose, glycerol, and lactate, respectively. Minimal medium provided unconstrained ammonium, phosphate and sulphate with glucose import at 10 mmol gr^−1^h^−1^ and O_2_ at 2 mmol gr^−1^h^−1^.

### Buffering-potentiation protocol regarding biochemical noise variability

We followed a procedure grounded on the one presented by Wang and Zhang (Wang and Zhang, 2011) to simulate the noise in the flux of a reaction. Flux noise incorporates experimentally measured gene expression noise data(Newman et al., 2006) that largely excluded extrinsic noise (noise measured in YPD conditions) and approximates the metabolic network as a linear pathway of length *n* **(Table S1)**. For a fixed *n*, we run 10000 simulations in which we constrain fluxes according to the noise and compute the corresponding fitness with MOMA (deviation to a noiseless metabolism) to obtain the variability associated to intrinsic noise (we presented *n* = 4 in the main text(Wang and Zhang, 2011)). We apply this procedure for each reference metabolism (wild-type and mutants) so that we can define a θ score for the variability in growth rate associated with noise: θ_noise_.

### Buffering-potentiation protocol regarding environmental variability

We generated 10000 different environments by randomly modifying the bounds of the nutrient reactions defining the YPD medium while maintaining ammonia, phosphate, sulphate, sodium, potassium, CO_2_, and H_2_O unbound and the import of O_2_ to 2 mmol gr^−1^h^−1^. Fitness of the new environmental conditions was computed with MOMA with the corresponding metabolic solution in YPD as reference and normalized by the fitness value of that reference metabolism. We computed the variability on this (relative) fitness to then define a θ score as before: θ_environment_.

### Pleiotropy and Epistasis

We applied FBA to compute the production rate of each biomass precursor for a given growth medium and genetic environment. To simulate the production of a given metabolite, we added a new exchange reaction to the model representing the secretion of this metabolite, and maximize the flux through this reaction (Shlomi et al., 2007; Szappanos et al., 2011). For the single-knockout annotation, we systematically deleted each gene and considered it as contributing to the production of a certain metabolite if its loss reduced the metabolite’s production rate more than 20%. We divided the number of metabolites for which a gene contributes by the total number to obtain a normalized score between 0 and 1. To compute the epistatic network, we calculated with FBA the growth rates of all single and double deletion mutants encompassing all nonessential genes. The mutant/ WT growth ratios obtained are used to compute an epistatic score (ε), which incorporates a multiplicative model and posterior scaling (Bajić et al., 2014; Segrè et al., 2005); interactions with |ε| < 0.01 were not considered.

### Phenotypic potential

Morphological phenotypes are available for two sets of nonessential (Levy and Siegal, 2008) and essential (Bauer et al., 2015) genes, in which a single measure of phenotypic variance termed the phenotypic potential was obtained. Note, however, that this measure is not completely equivalent for the two sets.

### Variation of expression of metabolic genes in cancer

We used a collection of 22 tumor types assembled by Hu et al. (Hu et al., 2013) to examine the variability in the expression of human metabolic genes in tumor and control samples. We specifically examined a list of metabolic genes obtained from the Human metabolic reconstruction model (Swainston et al., 2016). Each dataset was normalized using the RMA algorithm in the Bioconductor 3. 7 packages (www.bioconductor.org) running under R version 3.7. To obtain **Figure S1**, we quantified, for each enzyme, the standard deviation of expression in the corresponding control and tumor samples. We then calculated how many enzymes presented more variability in the tumor than in the control and divided that value by the entire number of enzymes to obtain a ratio. This defines the “ratio of enzymes with more variability in tumor conditions”. We also computed a null behavior for this ratio by randomization of expression values between control and tumor samples (to compute again the ratio, 1000 randomizations). Most tumor conditions led to an increase in expression variability.

---

**Figure S1.**
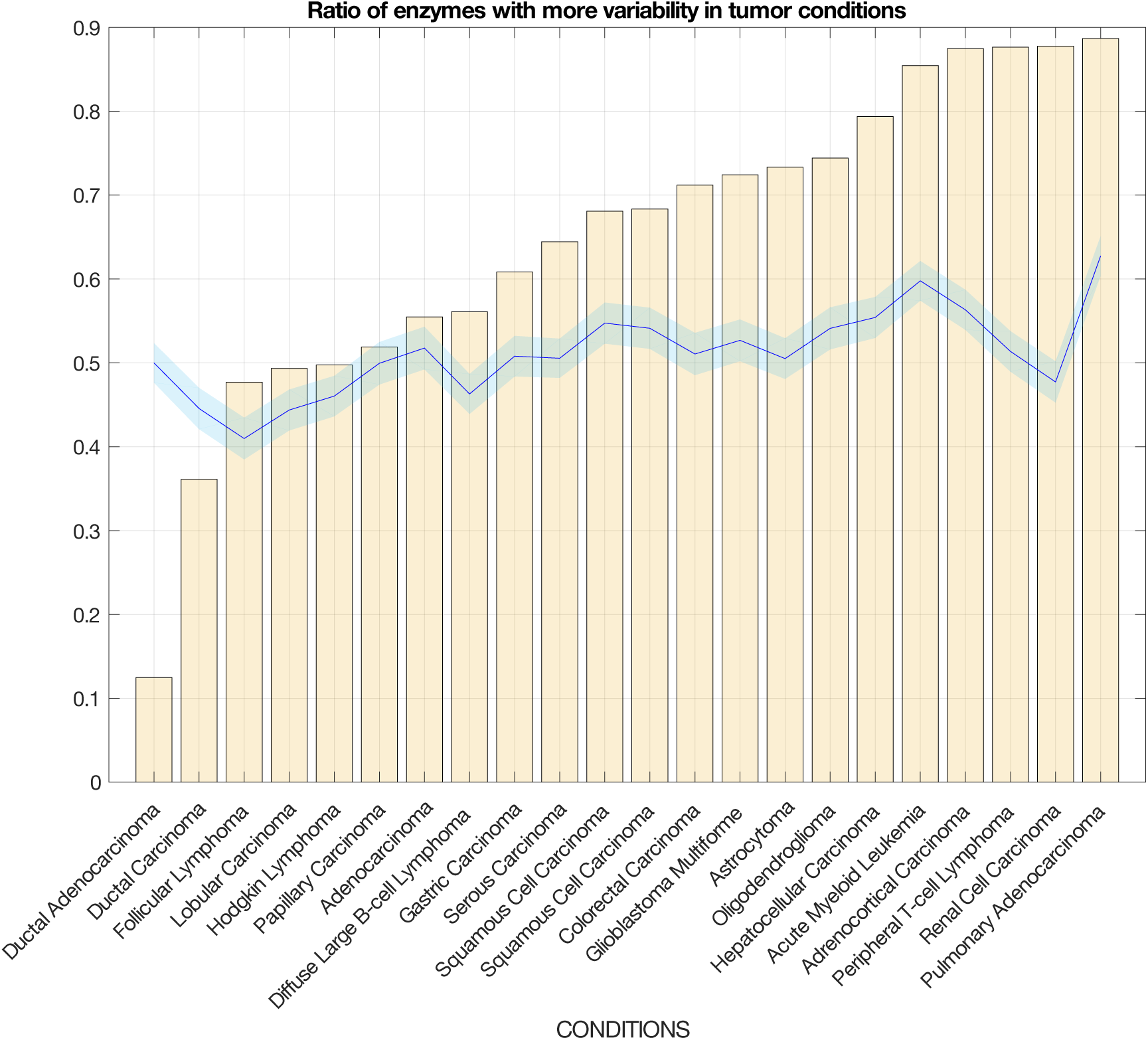
Mutations characterizing specific tumors increase the enzyme expression variability as compared to normal tissues. We used gene expression data of pairs of control and tumor samples to quantify the variability (standard deviation) in the expression of metabolic genes within each sample (see Methods for details). With these scores, we estimated the number of enzymes with more variation within the tumor sample than the control and then divided this value by the total number of enzymes considered (this ratio is indicated by the orange bars, sorted by increasing ratio). We also computed the expected null value of this ratio by randomization of expression data between tissue and control. We plot the mean null value of these randomizations (blue curve) and the +/-2 std (blue shading, 1000 randomizations).

